# Sensitivity of diffusion-tensor and correlated diffusion imaging to white-matter microstructural abnormalities: application in COVID-19

**DOI:** 10.1101/2022.09.29.510004

**Authors:** Nick Teller, Jordan A. Chad, Alexander Wong, Hayden Gunraj, Xiang Ji, Bradley J MacIntosh, Asaf Gilboa, Eugenie Roudaia, Allison Sekuler, Benjamin Lam, Chris Heyn, Sandra E Black, Simon J Graham, J. Jean Chen

## Abstract

There has been growing attention on the effect of COVID-19 on white-matter microstructure, especially among those that self-isolated after being infected. There is also immense scientific interest and potential clinical utility to evaluate the sensitivity of single-shell diffusion MRI methods for detecting such effects. In this work, the sensitivities of three single-shell-compatible diffusion MRI modeling methods are compared for detecting the effect of COVID-19, including diffusion-tensor imaging, diffusion-tensor decomposition of orthogonal moments and correlated diffusion imaging. Imaging was performed on self-isolated patients at baseline and 3-month follow-up, along with age- and sex-matched controls. We demonstrate through simulations and experimental data that correlated diffusion imaging is associated with far greater sensitivity, being the only one of the three single-shell methods to demonstrate COVID-19-related brain effects. Results suggest less restricted diffusion in the frontal lobe in COVID-19 patients, but also more restricted diffusion in the cerebellar white matter, in agreement with several existing studies highlighting the vulnerability of the cerebellum to COVID-19 infection. These results, taken together with the simulation results, suggest that a significant proportion of COVID-19 related white-matter microstructural pathology manifests as a change in water diffusivity. Interestingly, different b-values also confer different sensitivities to the effects. No significant difference was observed in patients at the 3-month follow-up, likely due to the limited size of the follow-up cohort. To summarize, correlated diffusion imaging is shown to be a sensitive single-shell diffusion analysis approach that allows us to uncover opposing patterns of diffusion changes in the frontal and cerebellar regions of COVID-19 patients, suggesting the two regions react differently to viral infection.

## Introduction

It is increasingly recognized that a major cause of death in COVID-19 is infection of the central-nervous system (CNS) (Iadecola et al., 2020). On the strength of earlier evidence from SARS and MERS, the nasal passage and blood-brain barrier serve as the main points of entry into the brain (Krasemann et al., 2022; Meinhardt et al., 2021). The regions surrounding the olfactory cortex, including contrast to noise ratio the orbitofrontal cortex, may be affected as suggested by reports of loss of olfaction (anosmia) in some COVID-19 patients with early variants (ENT UK at The Royal College of Surgeons of England, 2020).

Neurological effects of COVID are important considerations among those that self-isolated, but remain under-investigated, particularly the more subtle microstructural brain effects that are not visible on conventional clinical scans. Even mild COVID-19 can induce CNS damage (Fernández-Castañeda et al., 2022). Moreover, in an increasing portion of infected individuals, COVID-19 neurological and psychiatric effects may be prolonged substantially beyond the period of infection (i.e. the post-COVID condition) (Callard and Perego, 2020; Stefanou et al., 2022). There is, nonetheless, limited understanding of the pathophysiological mechanisms of the post-COVID condition.

Neuroinflammation has been suggested as the core pathophysiological change associated with viral infections of the CNS in both acute COVID-19 and the post-COVID stage (Lee et al., 2022; Shankar et al., 2008). The inflammatory response can lead to recruitment of immune cells and edema, and if unchecked, will lead to cell death. A hallmark of the early inflammatory process is the destruction of the myelin sheaths protecting the cerebral white matter (demyelination), which can be a promising search target. As the degree of neuroinflammation depends on the context, duration, and course of the primary viral invasion (DiSabato et al., 2016), so does the manifestation of inflammation. To this point, different types of neuroimaging tools are useful for capturing the early-acute, late-acute and subacute phases of infection. For instance, neuronal death can be observed as edema, ischemic lesions and tissue loss by macrostructural T1 and T2 magnetic resonance imaging (MRI) (Russo and McGavern, 2015), particularly in the white matter bordering grey-matter (Gilden, 2008). Conversely, white-matter microstructural changes are an earlier marker of neuroinflammation that can be captured using diffusion MRI. Diffusion MRI markers have been well correlated with disruptions in the blood-brain barrier and with biomarkers of infection and inflammation in the cerebrospinal fluid (Wright et al., 2015). Conventional diffusion tensor imaging (DTI) metrics describe the size and shape of the diffusion tensor (Pierpaoli and Basser, 1996), and typically can be derived from single-shell diffusion MRI acquisitions, which are the dominant type of acquisition in clinical research. The DTI method of acquisition and analysis provides estimates of mean diffusivity (MD) and fractional anisotropy (FA), but the latter is confounded by concurrent MD effects. The DT-DOME (orthogonal decomposition based on the eigenvalue moments) method was proposed as a simple solution to this challenge, whereby the diffusion pattern is characterized by the norm of anisotropy (NA) and mode of anisotropy (MO) instead of FA (Chad et al., 2021).

There has been a recent increase in studies that assess brain microstructure in COVID-19 (Esposito et al., 2022; Lu et al., 2020a). Notably, one was conducted amongst 60 hospitalized patients at baseline and 3-month follow-up (Lu et al., 2020a); reduced MD and axial diffusivity (AD) were found in the superior fronto-occipital fasciculus, the external capsule and the corona radiata. These observations were attributed to the accumulation of necrotic debris following neuroinflammation that impedes extracellular diffusion (Bhatt et al., 2017; Westman et al., 2019). The other study, focusing on a 1-year follow-up of hospitalized COVID-19 patients (Huang et al., 2021; Lu et al., 2020b), despite the absence of any significant patient-control difference in conventional DTI and diffusional-kurtosis parameters.

The literature presents several knowledge gaps: (1) both of these diffusion MRI studies were conducted on hospitalized patients, whereas the majority of patients are not being hospitalized during the pandemic, are self-isolated while infectious, and have not been well studied by medical imaging modalities; (2) these two studies disagree on the nature of microstructural effects of COVID-19, possibly as they were conducted at different follow-up times; (3) the lack of evidence regarding the post-COVID condition hamper our efforts to trace possible underlying disease mechanisms; (4) whereas one of the two studies showed COVID-related effects only when using multi-compartmental analysis of multi-shell data, single-shell DTI acquisitions are more practical clinically, and assessing the sensitivity of single-shell based metrics to COVID effects is valuable to the research community. It is thus likely that the sensitivity of existing single-shell metrics is insufficient, and it is beneficial to broaden the exploration of single-shell methods. One potential alternative presented in this work is a single-shell compatible method called correlated diffusion imaging (CDI) was proposed to enhance the difference between normal and cancerous prostate tissue (Wong et al., 2013). Given its sensitivity to prostate cancer, it would be useful to explore its utility in imaging COVID-19 effects. Thus, in this study, we aim to compare the sensitivity of single-shell DTI, DT-DOME and CDI in their abilities to address the above research gaps related to the imaging of COVID-19 effects on white matter microstructure.

## Methods

### Simulations

The theory of DTI, DT-DOME and CDI are detailed in the Theory section of the Supplementary Materials. Briefly, DT-DOME uses orthogonal tensor decompositon to generate unbiased parameters of anisotropy NA and MO (Chad et al., 2021), while CDI provides a marker based on multiplicative diffusion-image intensities that is sensitized to restricted diffusion (Wong et al., 2013). To better understand the relationship between conventional DTI, DT-DOME and CDI parameters and their sensitivities to pathology, and to demonstrate the application of the CDI concept to conventional brain diffusion imaging protocols, we simulated a set of diffusion signals arising from 34 diffusion directions, with a single b value of 700 s/mm^2^, plus a single b=0 volume. The signal intensity is simulated for two types of ground truth:

1. healthy white matter, with anisotropic diffusion (NA = 1 mm^2^/s), and an MD of 0.6 mm^2^/s;
2. diseased white matter, with a range of MD values from 80% to 120% of healthy tissue, also for an isotropic and anisotropic case.

We generated diffusion-weighted signals using Eq. 1, and added multiple instances of Rician noise, with the noisy signal defined as

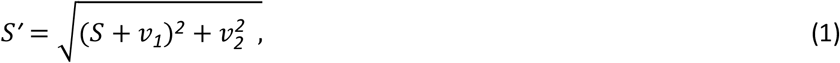

where *v*_*1*_and *v*_2_ are the real and imaginary components of the noise signal, both normally distributed. The SNR, defined as 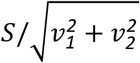, is set to vary from 10 to 110, and signals were obtained using 100 independent instances of Rician noise for each SNR, as is typical of diffusion images at 3 Tesla.

Since the CDI intensity is based on a product, it can quickly exhaust our data range. To compress the dynamic range of CDI values, a log transformation is applied to the product in Eq. A7 (Supplementary Materials) to provide the final CDI value (log(CDI)). A typical log(CDI) map generated from in vivo DTI data is shown in **Fig. A4** in Supplementary Materials. We further defined a range of fractional MD deviation of the simulated tissue from “healthy” (i.e. ground-truth MD of disease tissue)/(ground-truth MD of healthy tissue). We assessed the following metrics for comparing DTI and CDI performance over this range:

1. Effect size = {(estimated parameter for disease tissue) -(estimated parameter for healthy tissue)}/(estimated parameter for healthy tissue);
2. Contrast-to-noise ratio (CNR) = mean(effect size) / std(effect size) across multiple noise instances and SNRs.

### Study Participants

Participants in the current study were recruited between May 2020 and September 2021 through the Department of Emergency Medicine at Sunnybrook Health Sciences Centre, Toronto, Canada, physician referral, and community advertisements. Eligibility and consenting procedures were performed over phone or email. The Research Ethics Board at Sunnybrook Health Sciences Centre approved this study. Inclusion criteria for this study included being between 20 and 75 years of age and having documented evidence of a positive or negative COVID-19 diagnosis, as determined by a provincially approved facility through a nasopharyngeal or oropharyngeal swab and subsequent real-time reverse transcription polymerase chain reaction (RT-PCR) test. Exclusion criteria for this study included previous diagnosis of dementia, an existing neurological disorder, severe psychiatric illness, previous traumatic brain injury, on-going unstable cardiovascular disease, or contraindications to MRI (e.g., ferromagnetic implants).

### Study design

The current study stems from the NeuroCOVID-19 observational neuroimaging protocol (MacIntosh et al., 2021). We report on participants who were recruited to one of two groups: 1)COVID+ (N= 39) individuals that had tested positive for COVID-19 at an Ontario-approved facility, went into home isolation and have recovered from the acute symptoms; 2) COVID-(N=14) individuals who experienced flu-like symptoms but tested negative for COVID-19. There was no statistical difference between the mean age of the COVID+ and COVID-groups (see Table 1). 19 of the COVID+ participants were imaged at 3 months following the baseline imaging session.

**Table 1.** Patient demographics.

| Category | Patients (Initial)<br>(STDEV) | Patients (Follow-up)<br>(STDEV) | Controls<br>(STDEV) |
| --- | --- | --- | --- |
| n | 39 | 19 | 14 |
| Age | 42.1 | 45.3 | 43 |
| Sex | M:11;F:28 | M:9;F:10 | M:5;F:9 |
| Crystallized Composite Score | 100.3 (13.0) | 106.2 (13.0) | 107.9 (9.8) |
| Fluid Composite Score | 104.5 (16.1) | 109.6 (18.2) | 99.5 (17.5) |
| Negative Affect Summary | 60.4 (9.5) | 56.5 (8.1) | 58.1 (14.0) |
| Well Being Summary | 45.8 (11.6) | 46.7 (7.8) | 42.8 (8.8) |
| Social Satisfaction Summary | 48.1 (11.5) | 46.0 (9.5) | 39.9 (12.7) |

### Cognitive and Emotional Scores

The assessments are detailed in our recent publication (MacIntosh et al., 2021). Briefly, the NIH Toolbox full Cognition and Emotional Battery as well as the PROMIS tools (The Patient Reported Outcomes Measurement Information System, PROMIS-Canada) were administered using an iPad app, yielding Crystalized Intelligence Composite Scores, Fluid Intelligence Composite Scors, Negative Affect Memory, Well-being Summary and Social Satisfaction Summary Scores. Scores are summarized in **Table 1**.

### Image Acquisition

Diffusion MRI data were acquired on a Siemens Prisma 3T system with 5 b=0, 34 directions at b=700s/mm^2^, 34 directions at b=1400 s/mm^2^ and 34 directions at b=2100 s/mm^2^, with TR = 4.3 s, TE= 62ms, matrix size=96×96×60 (2×2×2 mm^3^ voxel resolution), 2-fold through-plane simultaneous multi-slice acceleration with 2 field-of-view shifts. A 1mm isotropic T1 anatomical image was also acquired for each participant, with TR/TE/TI = 2500/4.7/1100 ms, flip angle = 7°, FOV = 256×256×192 mm.

### Data Processing

All data sets were manually checked for quality. Data were corrected for eddy currents and susceptibility-related distortions using EDDY and TOPUP, respectively. FA, MD, AD, RD, NA and MO were obtained using the single-compartment DTI-model fit implemented through FMRIB Software Library (FSL 5.0) dtifit ((Jenkinson et al., 2012), using the b=0 and 700 s/mm^2^ shells. Correlated diffusion imaging (CDI) maps were computed based on the log transform of Eq. A9 with an in house MATLAB script, using data from all diffusion directions in each of the b = 0, 700, 1400, and 2100 s/mm^2^ shells. A CDI map was also generated for each b value separately. These resulted in FA, MD, AD, RD, NA, MO and CDI maps for all participants.

### Statistical Analysis

FSL’s Tract-Based Spatial Statistics (TBSS) were used for voxelwise statistical analysis of the FA, MD, AD, RD, NA, MO, and CDI data. Voxelwise differences for each parameter were assessed for patients versus controls, as well as between initial visit and follow up. Each map was then subjected to significance testing using FSL randomise with 500 permutations using threshold-free cluster en-hancement. Region of interest (ROI) analyses were performed on regions of significance from the voxelwise comparison. Group differences in cognitive and emotional scores were assessed using unpaired t-tests, thresholded at the 0.05 significance level. Cognitive and emotional scores with significant group differences were used as correlates in FreeSurfer’s General linear model (GLM) analysis to assess correlation with DTI parameters.

## Results

### Simulations

The simulations revealed an inverse relationship between CDI and MD values. As shown in **Fig. A1** in Supplementary Materials, increasing MD (mm^2^/s) is reflected in decreasing CDI, shown as increasingly negative log values (in this case for a SNR of 110). This is the case for both isotropic and anisotropic diffusion. However, no such relationship is discernible for CDI versus FA.

**Figure 1.**
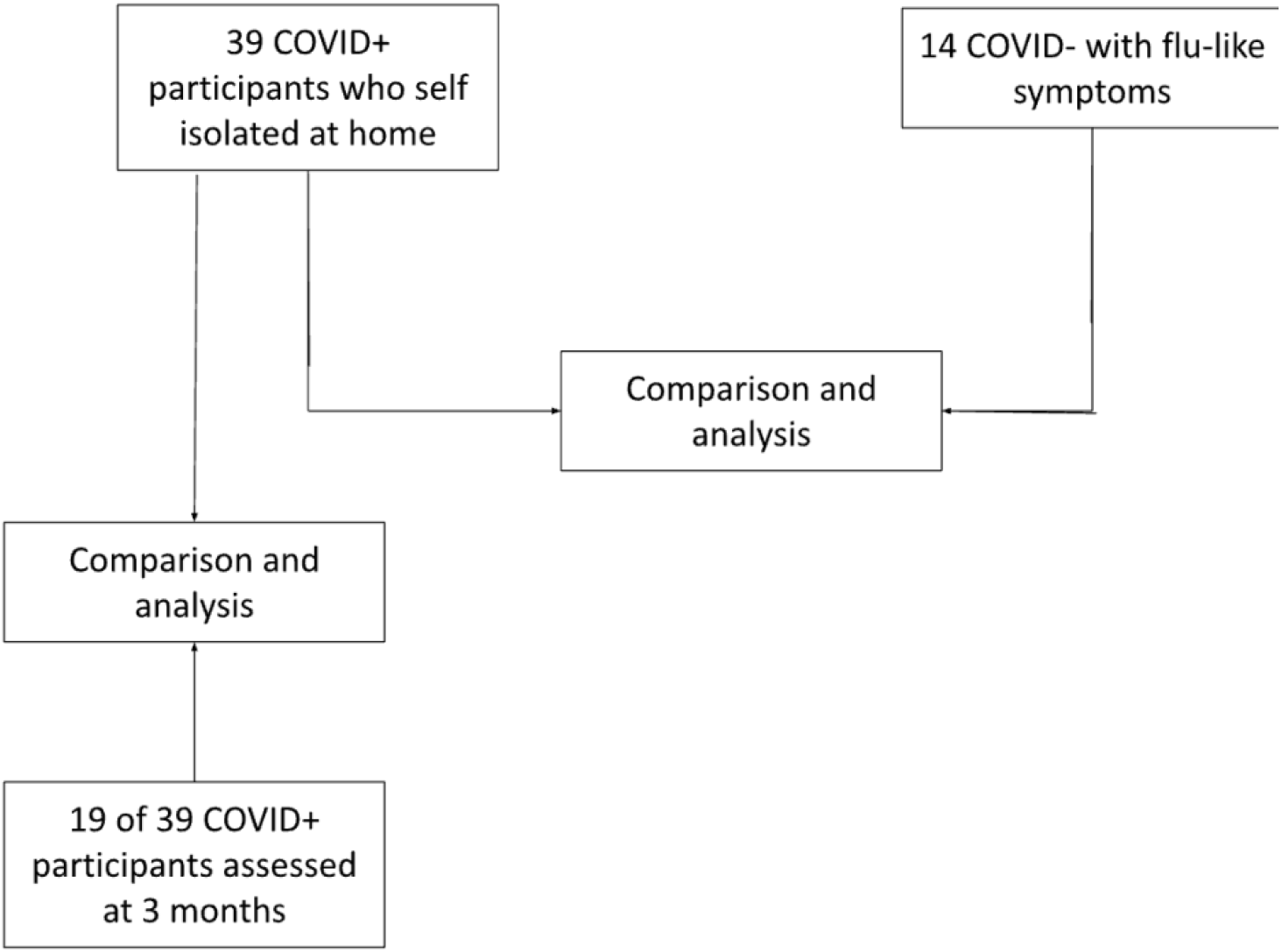
Flow chart of study design. Two sets of comparison were performed: (i) between COVID+ and COVID-groups; (ii) between the baseline and 3-month time points in the COVID+ group.

Shown in **Fig. 2**, are the estimated log(CDI) and corresponding MD, FA, NA and MO values for healthy and diseased tissue, averaged across all SNR and noise instances. When diseased tissue is modeled as variations in MD alone, increasing MD corresponds to decreasing log(CDI) and FA values, most consistently in anisotropic diffusion (**Fig. 2b**). The distinction between healthy and diseased tissue is unclear when viewed through NA and MO (**Fig. d, e)**. The effect sizes, i.e. percent differences between healthy and diseased tissue, computed for the estimated MD, FA and corresponding log(CDI) values, and averaged across all SNR and noise instances, are shown in **Fig. A2** of the Supplementary Materials. The log(CDI) and MD exhibit similar effect sizes, much higher than that of FA, NA and MO.

**Figure 2.**
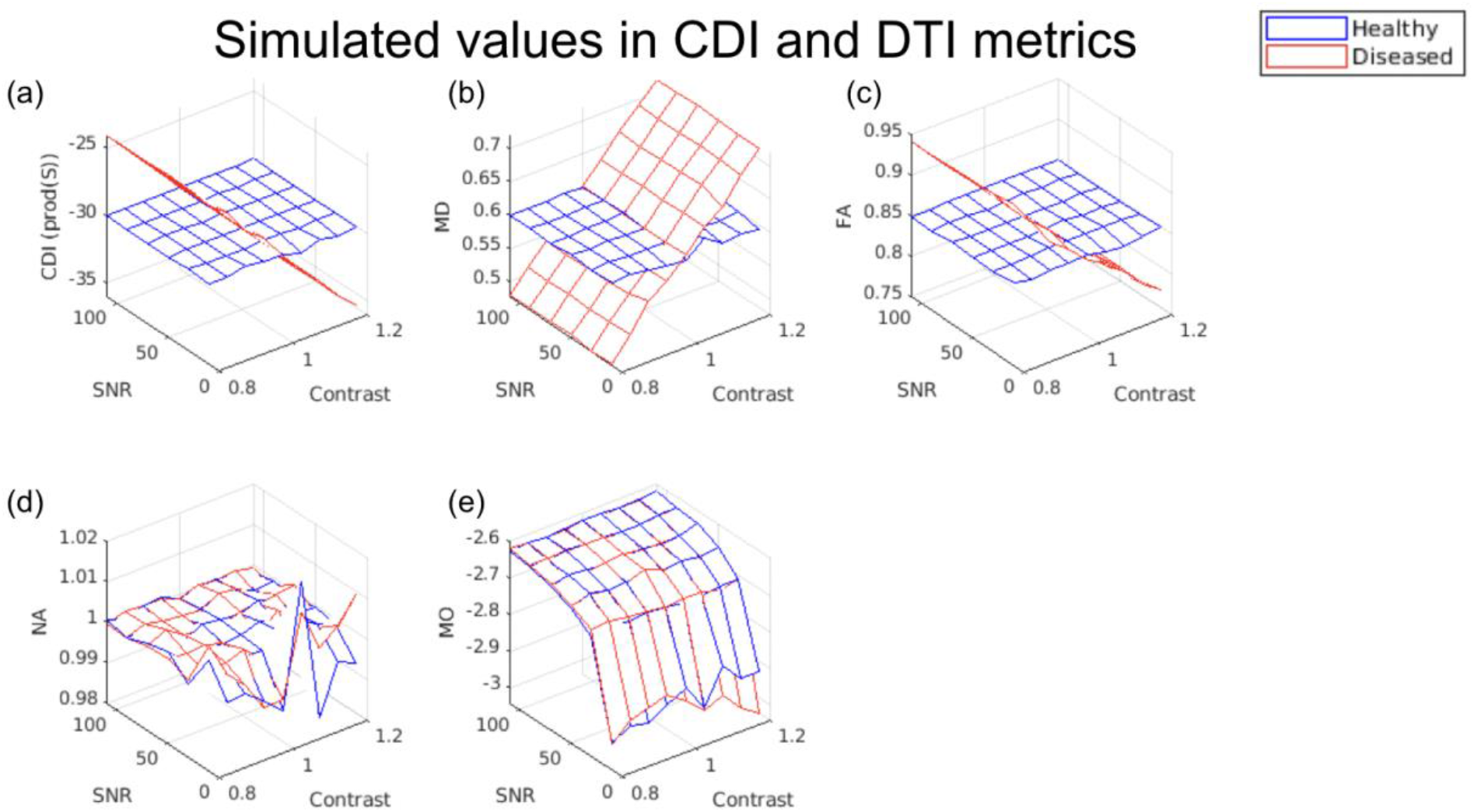
Estimated CDI and DTI metrics for healthy and diseased white-matter tissue. Each vertex on the 3D mesh represents 1 data point, computed as the average across all noise instances at each SNR. In all cases, a contrast value > 1 indicates higher MD in diseased tissue. Increasing MD (b) is reflected as decreasing log(CDI) for diseased tissue (a), with FA showing minimal difference between health and disease (c). Note that the difference between the simulated healthy and diseased tissues is in MD alone, so as expected, NA (d) and MO (e) exhibit negligible contrast between tissue types.

Shown in **Fig. 3** are the CNR comparisons for log(CDI), MD, FA, NA and MO. As CNR is based on both effect size and noise-related variability (encoded by the colour bar), a higher CNR is a robust indication of superior sensitivity to disease effects. log(CDI) is associated with maximum CNRs of 40 and higher (**Fig. 3a)** while MD and FA exhibit maximum CNRs of only 5-20 (**Fig. 3b,c)**. Lastly, NA and MO exhibit negligible CNR **(Fig. 3d, e)**.

**Figure 3.**
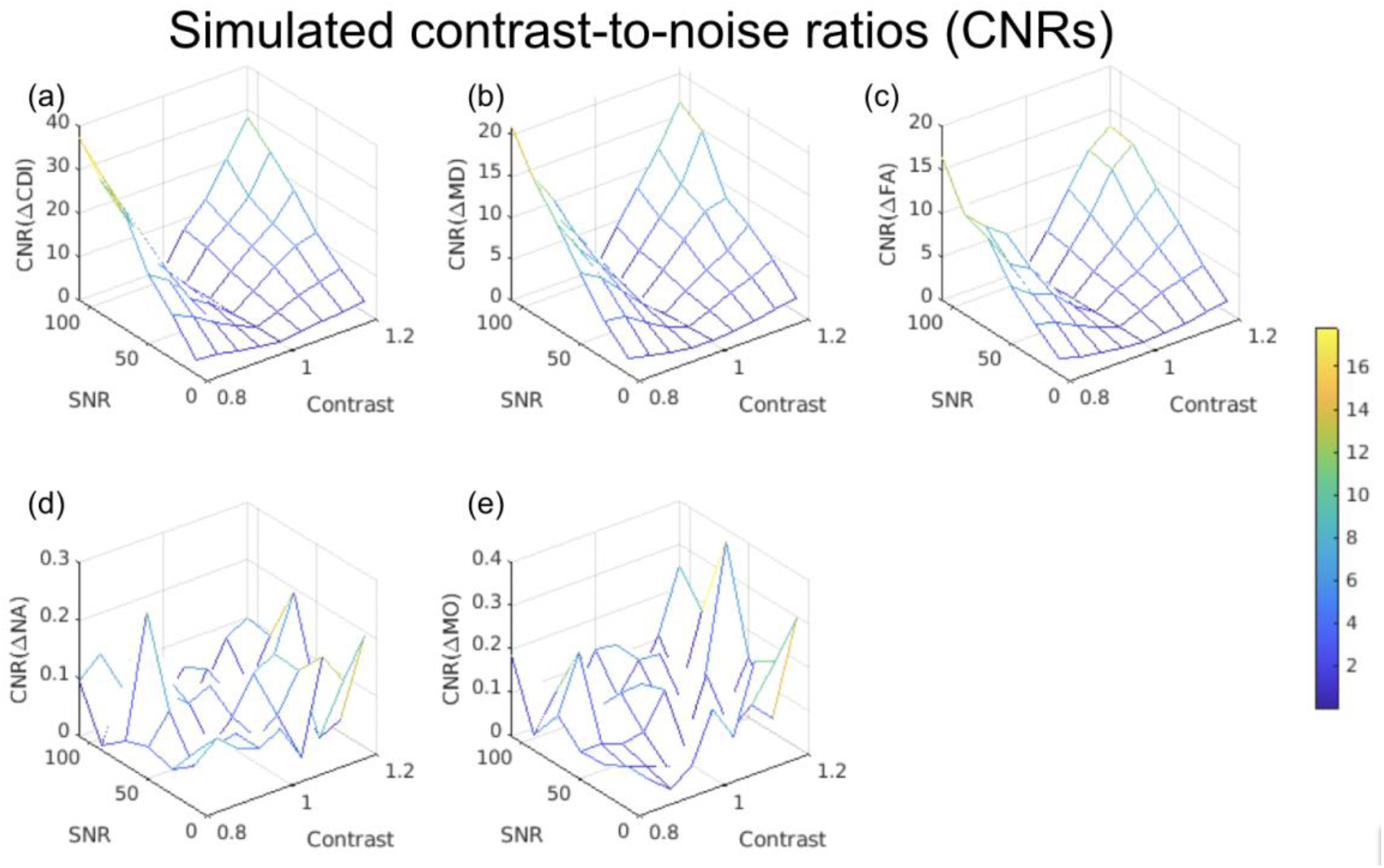
Estimated CNRs associated with CDI and DTI metrics for distinguishing healthy and diseased white-matter tissue. Each vertex on the 3D mesh represents 1 data point. The CNR represents the effect size normalized by estimation variability, and is encoded by the colour bar. Note that the difference between the simulated healthy and diseased tissues is in MD alone, so as expected, NA (d) and MO (e) exhibit negligible contrast between tissue types.

### Experimental results

We observed no significant difference between the baseline COVID+ and COVID-groups in terms of DTI metrics, including MD, FA, NA, AD, RD, and MO. However, we observed significant group differences through CDI values. Regions showing significantly higher log(CDI) in the COVID-group are shown in **Fig. 4**. The biggest log(CDI) differences between groups are encoded by the lowest b values. The b=1400 analysis reveals additional significant regions relative to the b=700 s/mm^2^ results, such as the genu of the corpus callosum. The b=2100 s/mm^2^ analysis shows least significance, although the superior corona radiata remains significant. **Fig. 5** shows boxplots of DTI metrics averaged over a union of these CDI ROIs of significant difference. High significance is observed in the corona radiata, with greatest significance in the age-controlled analysis. High significance is also evident in superior longitudinal fasciculus in all three analyses. Regions of high significance are consistent with literature (Huang et al., 2021), however with widespread frontal effects. **Fig. 5** demonstrates a general lack of significant group differences in these regions exhibited by conventional DTI and DT-DOME metrics.

**Figure 4.**
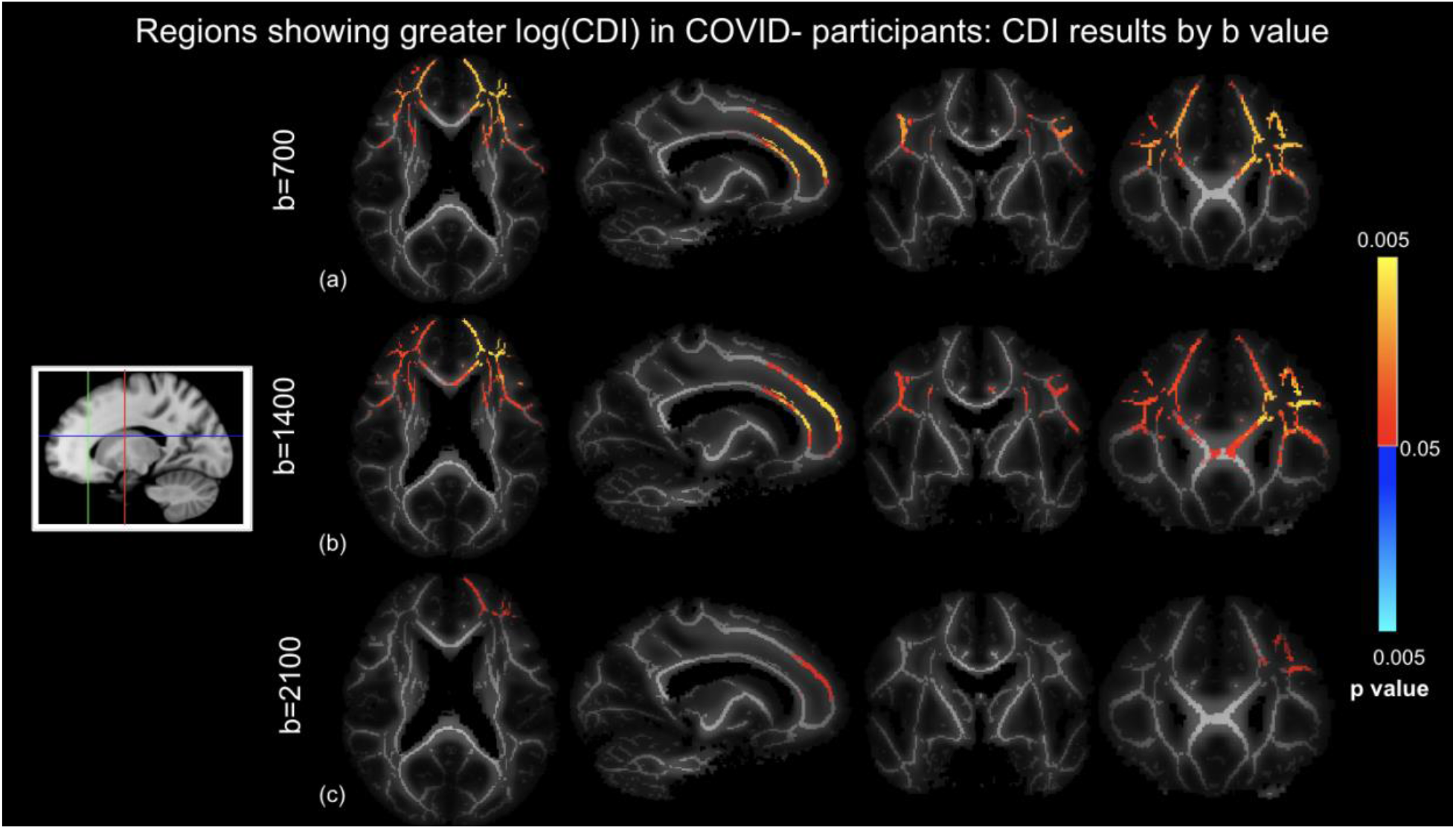
CDI comparison of COVID+ and COVID-groups based on different b values controlled for age and sex differences, where log(CDI) is greater in COVID-participants. (a) b=700 s/mm^2^, (a) b=1400 s/mm^2^, and (c) b = 2100 s/mm^2^.Orange-yellow indicate regions of statistically significant difference, with yellow indicating greater significance. Highest significance in the b=700 s/mm^2^ analysis. Slices are taken as shown in the icon to the left.

**Figure 5.**
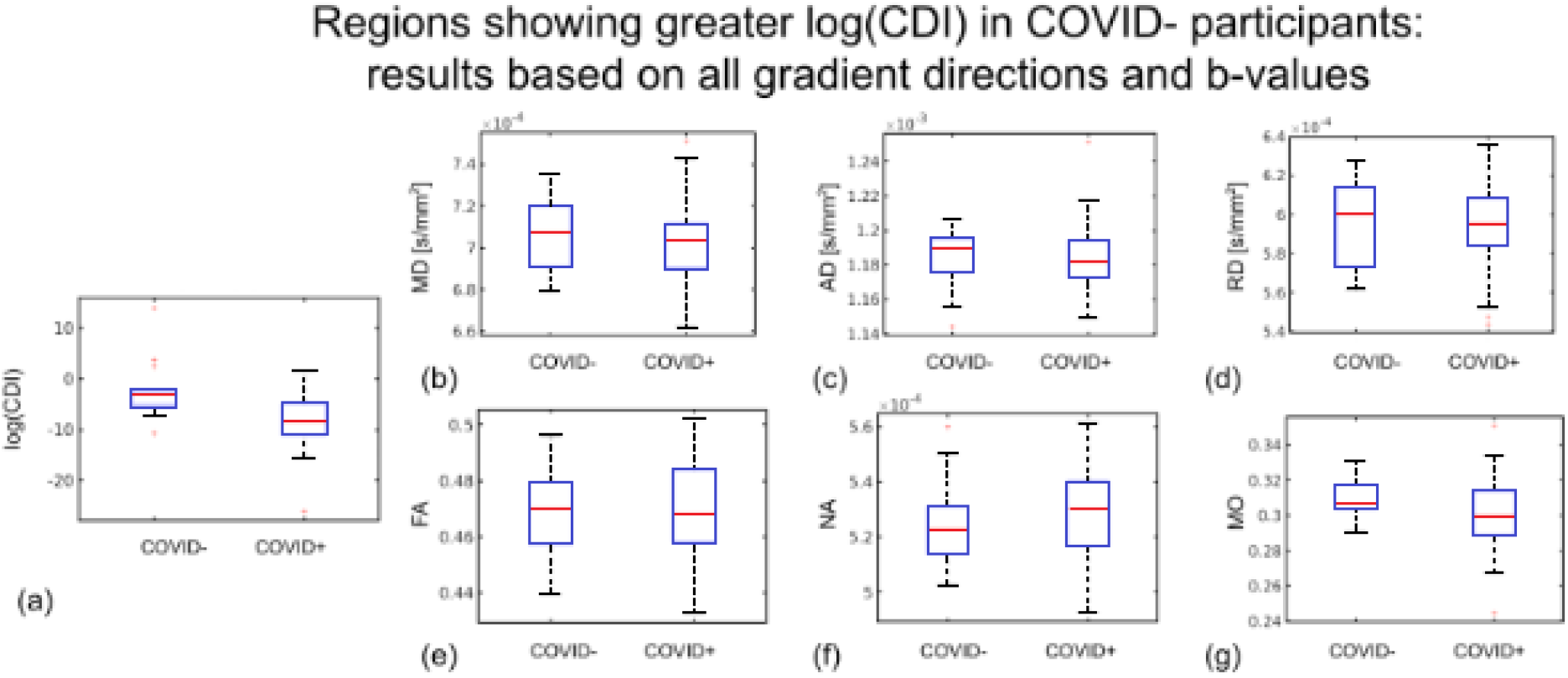
Region of interest (ROI) plots of DTI and log(CDI) values in regions where log(CDI) is greater in COVID-participants. The ROI is defined as TBSS analysis regions where log(CDI) was greater in COVID-participants based on all gradient directions and b-values in the TBSS analysis. Red line represents the median, the blue box represents the interquartile range, black lines represent minimum and maximum. Left boxes represent COVID-group and right boxes represent COVID+ group. Larger median in the COVID-group is evident in log(CDI) (a), MD (b), AD (c), RD (d), FA (e), and MO (g). Larger median in the COVID+ group is evident in NA (f).

Regions showing significantly higher log(CDI) in the COVID+ group are shown in **Fig. 6**. The biggest log(CDI) differences between groups are encoded by the highest b value. The b=2100 s/mm^2^ analysis reveals highest significance in the greatest number of voxels, particularly in the cerebellum. The b=700 s/mm^2^ analysis also reveals significant differences in the cerebellum. The b=1400 s/mm^2^ analysis reveals no significant differences. **Fig. 7** shows boxplots of DTI metrics averaged over a union of these CDI ROIs of significant difference. As seen in **Fig. 6**, high significance is evident in the cerebellum. **Fig. 7** demonstrates a general lack of significant group differences in these regions exhibited by conventional DTI and DT-DOME metrics.

**Figure 6.**
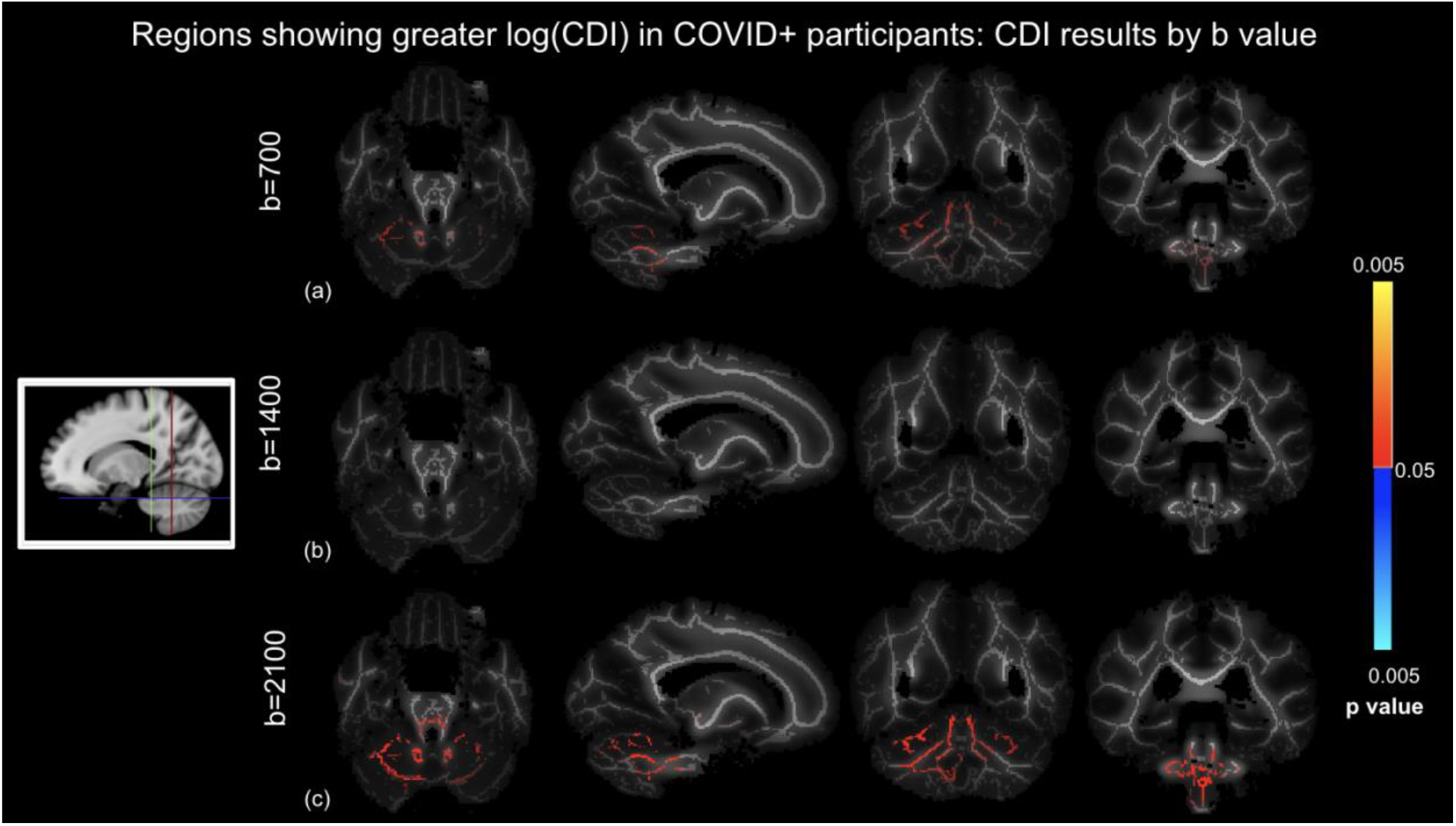
CDI comparison of COVID+ and COVID-groups using different b values controlled for age and sex differences, where log(CDI) is greater in COVID+ participants. (a) b=700 s/mm^2^, (b) b=1400 s/mm^2^, and (c) b = 2100 s/mm^2^. Red regions are statistically significant. Highest significance in the b=2100 s/mm^2^ analysis. Slices are taken as shown in the icon to the left.

**Figure 7.**
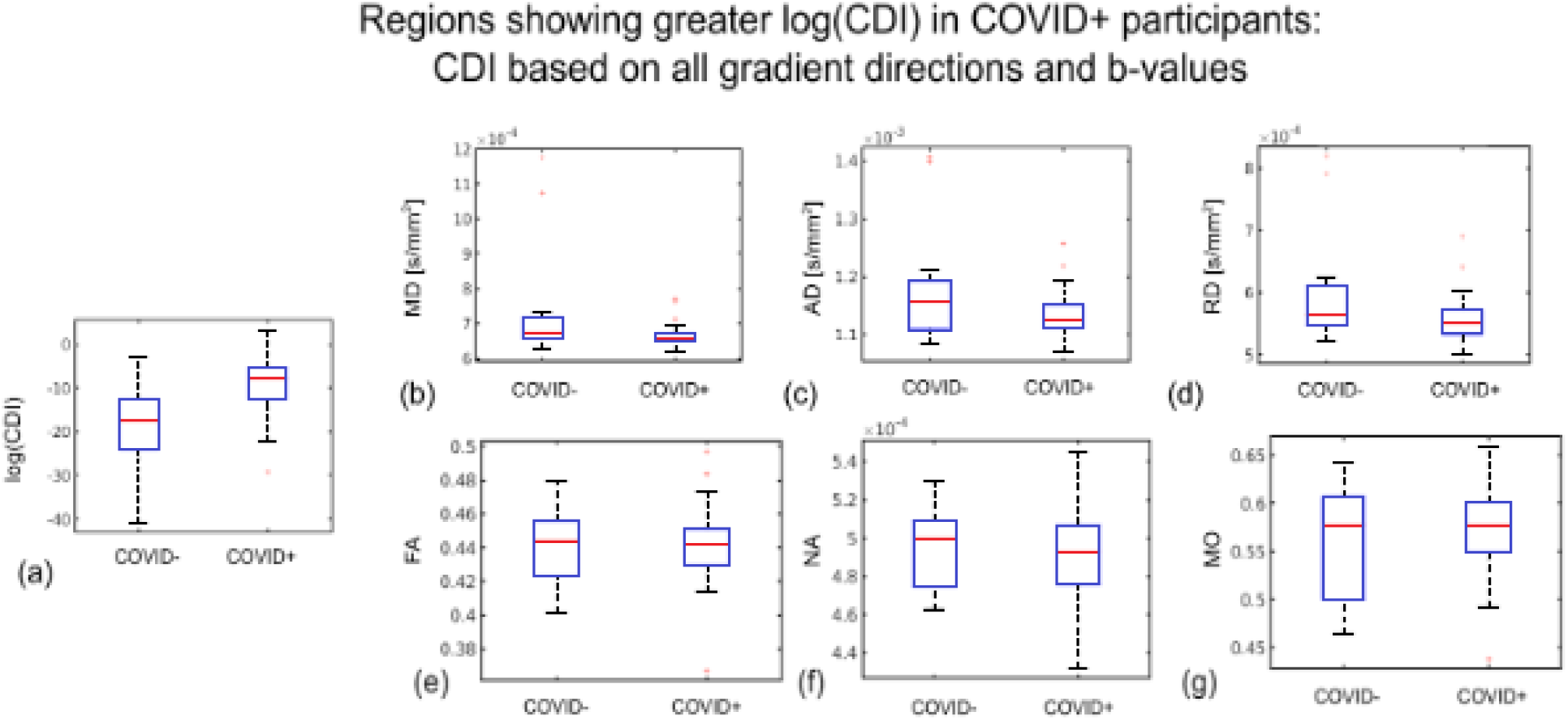
Region of interest (ROI) plots of DTI and log(CDI) values in regions where log(CDI) is greater in COVID+ participants. The ROI is defined as regions where log(CDI) was greater in COVID+ participants based on all gradient directions and b-values in the TBSS analysis. Red line represents the median, the blue box represents the interquartile range, black lines represent minimum and maximum. Left boxes represent COVID-group and the right boxes represent COVID+ group. Larger median in the COVID+ group is evident in log(CDI) (a) and MO (g). Larger median in the COVID-group is evident in MD (b), AD (c), RD (d), FA (e), and NA (f).

Lastly, in **Fig. 8**, we show the comparison of the log(CDI) results against recently published results using multi-compartmental diffusion modeling (NODDI). The three-compartment NODDI model revealed reduced intracellular water fraction (V_ic_) in the genu of the corpus callosum, the superior longitudinal fasciculus and the corona radiata, suggesting reduced axonal density in patients (Huang et al., 2021). Note that the regions of significant group difference in the current study are highly similar to those revealed by NODDI in a different population. Lastly, no significant differences were found between the initial and 3-month follow-up analysis, and expanded longitudinal evaluation is in progress.

**Figure 8.**
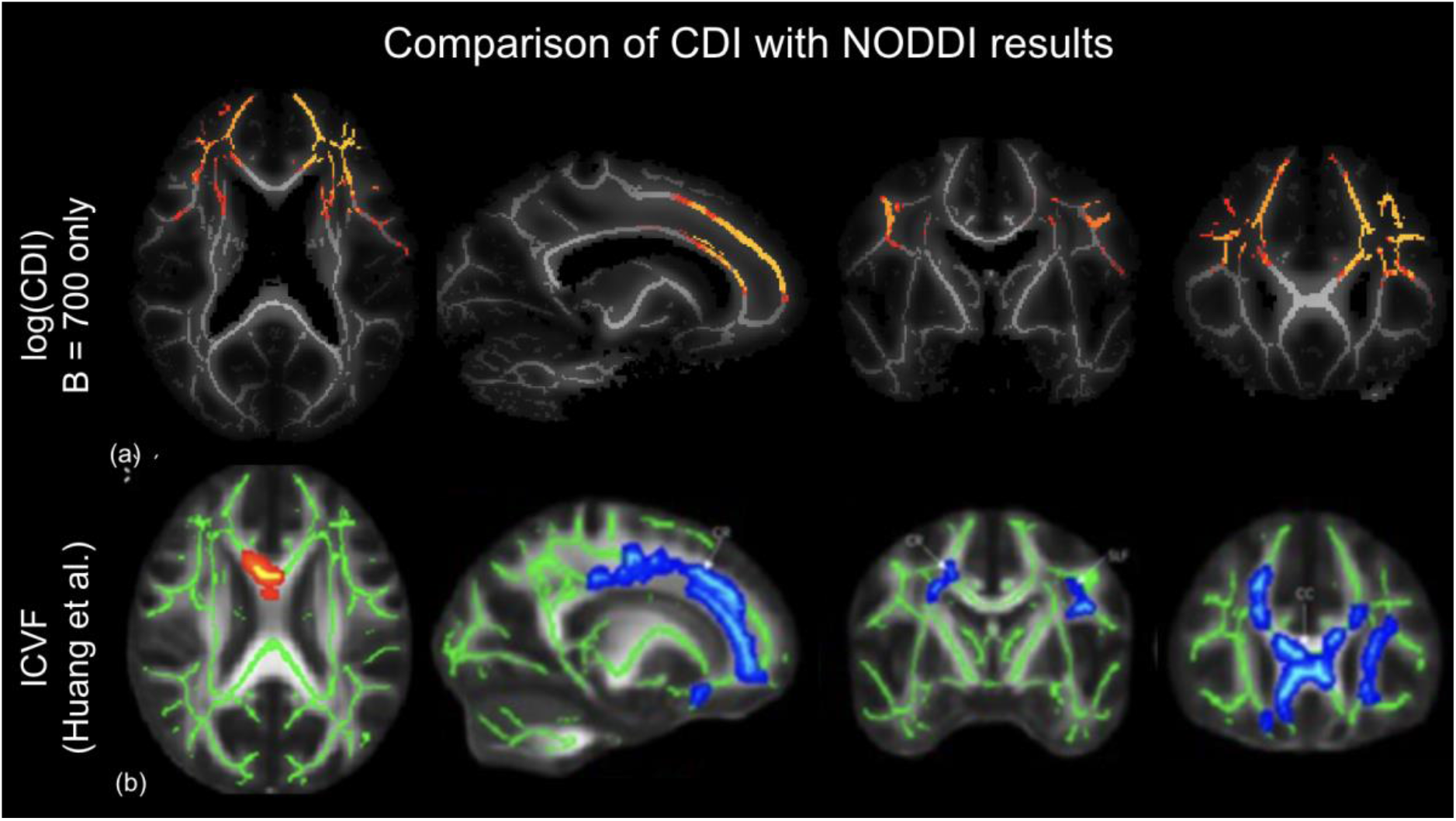
Comparison of current findings to findings in the literature. TBSS of log (CDI) results from the b=700 analysis (a) to ICVF results from NODDI analysis (b) (Huang et al., 2021). Orange-yellow regions in CDI analysis indicate regions of statistically significant higher log(CDI) in the COVID-group, with yellow indicating greater significance. Blue-light blue in the ICVF analysis indicates regions of higher V_ic_ in a control group than a COVID+ group with dark blue representing higher V_ic_ and light blue representing lower V_ic_. Both analyses reveal abnormal diffusion in the bilateral corona radiata of patients.

## Discussion

There are numerous well-documented impacts of COVID-19 on the brain in the literature. It is clear that COVID can not only enter the brain (Reiken et al., 2022), but can have numerous metabolic, immune, and hematologic effects (Shaikh et al., 2022). In this work, we pioneer the application of correlated diffusion imaging (CDI) in neuroimaging of COVID-19. We show that whereas DTI metrics did not reveal differences between the white matter of COVID- and COVID+ groups, log(CDI) exhibits highly significant differences. Compared to the previous instances of CDI applications (so far limited to prostate cancer) (Khalvati et al., 2016; Wong et al., 2021, 2015, 2013), our implementation of CDI involves 3 key differences: (i) instead of relying on images acquired at multiple gradient strengths, we rely mainly on the multiple diffusion directions, which are part of typical brain DTI acquisitions; (ii) given the small spatial scale of white-matter structures, we opted to not use a spatial probability distribution function in order to avoid spatial blurring and partial-volume effects; (iii) to compress the large dynamic range of brain CDI values and normalize the data distribution, we used log(CDI) in this work. Through simulations and experimental data, we demonstrate that CDI’s sensitivity to disease processes is independent of these processing choices. Based on prior literature in prostate cancer (Wong et al., 2021, 2013), a higher log(CDI) can be loosely interpreted as reflecting a denser tissue structure with restricted diffusion. Interestingly, we find 2 dichotomous trends in log(CDI) of COVID patients: (i) in anterior white matter, patients exhibit lower log(CDI) than controls, suggesting less restricted diffusion in patients; (ii) in the cerebellum, patients exhibit higher log(CDI) than controls, suggesting more restricted diffusion in patients.

### Simulations

In our simulations we assumed only single-shell diffusion data was available, and compared the sensitivities of DTI (MD, FA), DT-DOME (NA, MO) and CDI metrics through CNR. The simulated case covers a single-b-value scenario, with the number of diffusion directions (i.e. 34) identical to that of our MRI acquisitions. Disease conditions were assumed to range from reduced to enhanced diffusivity, relative to healthy tissue. Noise was added over a large range of SNRs, from 10 to 120, to cover a variety of image qualities. We demonstrated that in principle, CDI can be viewed as inversely proportional to MD, but that its sensitivity to disease is more than twice that of MD. Note that the disease effect is modeled as a difference in MD alone, so as expected, FA was less affected by the condition, whereas NA and MO do not exhibit contrast between healthy and disease, as they are not confounded by MD (unlike FA). Based on the simulated results, we hypothized that in cases where the pathology is defined by a change in diffusivity rather than anisotropy, the in vivo performance of CDI would be superior to that of DTI and DT-DOME.

### Potential sources of less restricted diffusion

Huang et al. acquired multi-shell diffusion MRI data with b values of 1000 and 2000 s/mm2, and showed that the NODDI parameter of intracellular water volume (Vic) alone demonstrated significant differences between patient and control groups. The reduced Vic in patients was interpreted as a reduction in axonal density following edema and axonal apoptosis (Huang et al., 2021). The affected areas include the superior longitudinal fasciculus, the genu of the corpus callosum and the bilateral corona radiata.

The strong anterior emphasis of these effects may be traced back to the role of the olfactory bulb (OB) as the most likely entry point for SARS-CoV-2 in the brain (Serrano et al., 2022; Xydakis et al., 2021). The OB has been shown to be the most likely brain region to contain SARS-CoV-2 and shows significant changes in gene expression in COVID-19 (Serrano et al., 2022). Moreover, studies examining the brain vasculature in deceased COVID-19 patients showed strong fibrinogen staining in the OB, indicating a leaky blood-brain barrier (BBB) at autopsy (Lee et al., 2022). Neurovascular studies demonstrated widespread BBB leakage and immune cell infiltration in the brains of COVID-19 patients (Lee et al., 2022). BBB breakdown in COVID-19 is characterized by fibrinogen accumulation in perivascular space (Lee et al., 2022; Wenzel and Schwaninger, 2022). Fibrinogen has been demonstrated to contribute to neuronal loss in animal models (Ryu and McLarnon, 2009). Fibrinogen has also been shown to activate microglia, the brain’s resident immune cells, in patients with Alzheimer’s disease (McLarnon, 2021). The products of microglial activation include pro-inflammatory cytokines and reactive oxygen species which contribute to inflammation and neurodegeneration (Akiyama et al., 2000). Thus, abnormalities in diffusion observed in this work may be a consequence of neuroinflammatory effects.

In our study the regions with significantly lower log(CDI) in patients align well with those identified by Huang et al. (Huang et al., 2021) (**Fig. 8**). As lower log(CDI) suggests less restricted diffusion, the interpretation also aligns with that of Huang et al. However, as shown in **Fig. 4**, multi-shell data is not necessary for uncovering this effect. In fact, the use of the b=700 s/mm^2^ shell alone yielded the most sensitive log(CDI) values. As the lowest b-value is sensitive to the fastest diffusion, and as CDI is boosted by restricted diffusion, this finding may indicate that the lower log(CDI) in patients is preferentially reflecting an enhancement of the fastest-moving water molecules, potentially reflecting enlargement of extracellular spaces (Huang et al., 2021) and/or leakage of the BBB. The literature indicates that the BBB is compromised early in the disease process, which also allows for viral effects to persist (Malerba et al., 2021). Although the precise mechanism of vascular damage remains hypothetical, it has been suggested that immunoglobulin complexes, cytokines, autoantibodies, and the SARS-CoV-2 virus itself may activate the endothelium and induce BBB leakage (Wenzel and Schwaninger, 2022).

### Potential sources of more restricted diffusion

For the first time, the current study reports diffusion abnormalities in the white matter of the cerebellum, where the higher log(CDI) values suggest more restricted diffusion. While this finding echoes findings by Lu et al. in anterior cerebral white matter (Lu et al., 2020a), it is opposite those of the anterior cerebral white matter found in the current study. Lu et al. reported generally reduced diffusivity, interpreted as the result of post-inflammatory debris accumulation (Lu et al., 2020a).

Several studies provide precedent for the impact of COVID-19 on the cerebellum. Notably, an MRI study from the UK Biobank discovered cerebellum volume loss in COVID patients (Douaud et al., 2022). Several case reports of COVID patients found acute cerebellitis following infection with SARS-CoV-2 (Fadakar et al., 2020; Moreno-Escobar et al., 2021). Furthermore, there is significant neuropathological evidence that COVID-19 enters the cerebellum, as several studies indicate that immune cell infiltration is most pronounced in the cerebellum (Colombo et al., 2021; Matschke et al., 2020). Moreover, one study examining biochemistry in the brains of COVID-19 patients found that the cerebellum in COVID-19 patients exhibits leaky Ca^2+^ channels and activity consistent with intracellular calcium leakage, which were suggested to contribute to the disease process (Reiken et al., 2022).

The BBB is compromised in the cerebellum of COVID-19 patients, as evidenced by fibrinogen infiltration in the perivascular space (Lee et al., 2022). Strong fibrinogen staining was evident in both the forebrain and cerebellum, however weak fibrinogen staining was only evident in the cerebellum (Lee et al., 2022). Furthermore, immune cell infiltration is most pronounced in the cerebellum (Colombo et al., 2021; Matschke et al., 2020) which may be indicative of susceptible vasculature. These findings are taken from post-mortem studies in individuals who died days or weeks after infection, indicating that viral entry in the OB and BBB leakage occurs early in the disease process. Our finding that log(CDI) is greater in patients in the cerebellum at the initial visit may be explained by debris accumulation following immune cell infiltration (Lee et al., 2022).

In our study we also uncovered previously unseen regions of significantly higher log(CDI) value in patients (**Fig. 10**), which suggests more restricted diffusion. As shown in **Fig. 6**, multi-shell data is not necessary for uncovering this effect, either. In fact, the use of the b=2100 s/mm^2^ shell alone yielded the greatest CDI sensitivity to this effect. As the highest b-value is sensitive to the slowest diffusion, and as CDI is boosted by restricted diffusion, this finding may indicate that the greater log(CDI) in patients is preferentially reflecting a restriction of the slowest-moving water molecules. Such molecules could be intra-cellular or myelinated water, and their restriction could reflect cytotoxic intra-cellular edema, which in turn can be caused by hypoxic insults in COVID-19 (van den Enden et al., 2020). As to why such effects are only seen in the cerebellum, this region consistently displays neuropathological abnormalities including immune cell infiltration which may be indicative of susceptible vasculature or increased viral entry (Matschke et al., 2020). Furthermore, while in the forebrain fibrinogen staining is localized around the damaged vessels, in the cerebellum fibrinogen staining is evident at greater distances from the vessels (Lee et al., 2022). Despite the OB being the most probable site of entry for COVID-19, pathological brain findings are consistently most prominent in the cerebellum. Immune cell infiltration, widespread fibrinogen leakage, and debris accumulation may be responsible for the restricted diffusion identified in this investigation.

### Limitations

This is the first instance of CDI technique being applied in neuroimaging, and we are cognizant of the uncertainties in our understanding of COVID-19. The fact that CDI was exceptionally sensitive to self-isolated COVID-19 white-matter pathology is encouraging, but also raises questions: (1) How does CDI perform inthe grey matter? (2) How does it reflect neurodegenerative processes in other conditions, such as aging and Alzheimer’s disease? (3) Under what circumstances would CDI fail to show sensitivity to pathology? These questions will propel our future studies to further validate and characterize this novel technique.

Another limitation of this investigation is that although DTI is sensitive to microstructure changes, it is not specific. For example, it is unclear whether, in the long term, restricted diffusion is an indicator of compensatory neurogenesis or persistent inflammation (Goldberg et al., 2021). To shed light on the interpretation of our findings, further research (potentially involving autopsy) is key.

We are also constrained by the limited longitudinal follow-up. Although short term COVID brain effects appear to manifest principally in the cortex and cerebellum, long term effects are more widespread. One area of the brain that becomes affected in long COVID is the cingulate cortex, which exhibited hypometabolism in patients with brain fog several months after contracting COVID (Hugon et al., 2022). The literature also indicates that the globus pallidum and substantia nigra show restricted diffusion at follow-up (Abdo et al., 2021). It is possible the sample size of the longitudinal follow-up was insufficiently powered to detect the changes observed in the literature. However, our longitudinal evaluations are ongoing, and our future work will involve a larger sample size, particularly in the initial visit vs follow-up analysis, to provide greater power to the analysis.

## Acknowledgments

We thank the CIHR, NSERC, Sandra Black Centre for Brain Resilience & Recovery, and the Sunnybrook Hospital Foundation for financial support.

## Notes

### Competing Interest Statement

The authors have declared no competing interest.

## References

Abdo, W.F., Broerse, C.I., Grady, B.P., Wertenbroek, A.A.A.C.M., Vijlbrief, O., Buise, M.P., Beukema, M., van der Kuil, M., Tuladhar, A.M., Meijer, F.J.A., van der Hoeven, J.G., 2021. Prolonged Unconsciousness Following Severe COVID-19. Neurology 96, e1437–e1442.

Akiyama, H., Arai, T., Kondo, H., Tanno, E., Haga, C., Ikeda, K., 2000. Cell Mediators of Inflammation in the Alzheimer Disease Brain. Alzheimer Disease and Associated Disorders. https://doi.org/10.1097/00002093-200000001-00008

Bhatt, N., Gupta, N., Soni, N., Hooda, K., Sapire, J.M., Kumar, Y., 2017. Role of diffusion-weighted imaging in head and neck lesions: Pictorial review. The Neuroradiology Journal. https://doi.org/10.1177/1971400917708582

Callard, F., Perego, E., 2020. How and why patients made Long Covid. Soc. Sci. Med. 113426.

Chad, J.A., Pasternak, O., Chen, J.J., 2021. Orthogonal Moment Diffusion Tensor Decomposition Reveals Age-Related Degeneration Patterns in Complex Fibre Architecture. Neurobiol. Aging. https://doi.org/10.1016/j.neurobiolaging.2020.12.020

Colombo, D., Falasca, L., Marchioni, L., Tammaro, A., Adebanjo, G.A.R., Ippolito, G., Zumla, A., Piacentini, M., Nardacci, R., Del Nonno, F., 2021. Neuropathology and Inflammatory Cell Characterization in 10 Autoptic COVID-19 Brains. Cells 10. https://doi.org/10.3390/cells10092262

DiSabato, D.J., Quan, N., Godbout, J.P., 2016. Neuroinflammation: the devil is in the details. Journal of Neurochemistry. https://doi.org/10.1111/jnc.13607

Douaud, G., Lee, S., Alfaro-Almagro, F., Arthofer, C., Wang, C., McCarthy, P., Lange, F., Andersson, J.L.R., Griffanti, L., Duff, E., Jbabdi, S., Taschler, B., Keating, P., Winkler, A.M., Collins, R., Matthews, P.M., Allen, N., Miller, K.L., Nichols, T.E., Smith, S.M., 2022. SARS-CoV-2 is associated with changes in brain structure in UK Biobank. Nature. https://doi.org/10.1038/s41586-022-04569-5

ENT UK at The Royal College of Surgeons of England, 2020. Loss of sense of smell as marker of COVID-19 infection. Royal College of Surgeons of England.

Esposito, F., Cirillo, M., De Micco, R., Caiazzo, G., Siciliano, M., Russo, A.G., Monari, C., Coppola, N., Tedeschi, G., Tessitore, A., 2022. Olfactory loss and brain connectivity after COVID-19. Hum. Brain Mapp. 43, 1548–1560.

Fadakar, N., Ghaemmaghami, S., Masoompour, S.M., Shirazi Yeganeh, B., Akbari, A., Hooshmandi, S., Ostovan, V.R., 2020. A First Case of Acute Cerebellitis Associated with Coronavirus Disease (COVID-19): a Case Report and Literature Review. Cerebellum 19, 911–914.

Fernández-Castañeda, A., Lu, P., Geraghty, A.C., Song, E., Lee, M.-H., Wood, J., Yalçın, B., Taylor, K.R., Dutton, S., Acosta-Alvarez, L., Ni, L., Contreras-Esquivel, D., Gehlhausen, J.R., Klein, J., Lucas, C., Mao, T., Silva, J., Peña-Hernández, M.A., Tabachnikova, A., Takahashi, T., Tabacof, L., Tosto-Mancuso, J., Breyman, E., Kontorovich, A., McCarthy, D., Quezado, M., Hefti, M., Perl, D., Folkerth, R., Putrino, D., Nath, A., Iwasaki, A., Monje, M., 2022. Mild respiratory SARS-CoV-2 infection can cause multi-lineage cellular dysregulation and myelin loss in the brain. bioRxiv. https://doi.org/10.1101/2022.01.07.475453

Gilden, D.H., 2008. Brain imaging abnormalities in CNS virus infections. Neurology 70, 84.

Goldberg, E., Podell, K., Sodickson, D.K., Fieremans, E., 2021. The brain after COVID-19: Compensatory neurogenesis or persistent neuroinflammation? EClinicalMedicine 31, 100684.

Huang, S., Zhou, Z., Yang, D., Zhao, W., Zeng, M., Xie, X., Du, Y., Jiang, Y., Zhou, X., Yang, W., Guo, H., Sun, H., Liu, P., Liu, J., Luo, H., Liu, J., 2021. Persistent white matter changes in recovered COVID-19 patients at the 1-year follow-up. Brain. https://doi.org/10.1093/brain/awab435

Hugon, J., Msika, E.-F., Queneau, M., Farid, K., Paquet, C., 2022. Long COVID: cognitive complaints (brain fog) and dysfunction of the cingulate cortex. J. Neurol. 269, 44–46.

Iadecola, C., Anrather, J., Kamel, H., 2020. Effects of COVID-19 on the Nervous System. Cell. https://doi.org/10.1016/j.cell.2020.08.028

Jenkinson, M., Beckmann, C.F., Behrens, T.E.J., Woolrich, M.W., Smith, S.M., 2012. FSL. Neuroimage 62, 782–790.

Khalvati, F., Zhang, J., Baig, S., Haider, M.A., Wong, A., 2016. Sparse Correlated Diffusion Imaging: A New Computational Diffusion MRI Modality for Prostate Cancer Detection. Journal of Computational Vision and Imaging Systems. https://doi.org/10.15353/vsnl.v2i1.107

Krasemann, S., Haferkamp, U., Pfefferle, S., Woo, M.S., Heinrich, F., Schweizer, M., Appelt-Menzel, A., Cubukova, A., Barenberg, J., Leu, J., Hartmann, K., Thies, E., Littau, J.L., Sepulveda-Falla, D., Zhang, L., Ton, K., Liang, Y., Matschke, J., Ricklefs, F., Sauvigny, T., Sperhake, J., Fitzek, A., Gerhartl, A., Brachner, A., Geiger, N., König, E.-M., Bodem, J., Franzenburg, S., Franke, A., Moese, S., Müller, F.-J., Geisslinger, G., Claussen, C., Kannt, A., Zaliani, A., Gribbon, P., Ondruschka, B., Neuhaus, W., Friese, M.A., Glatzel, M., Pless, O., 2022. The blood-brain barrier is dysregulated in COVID-19 and serves as a CNS entry route for SARS-CoV-2. Stem Cell Reports 17, 307–320.

Lee, M.H., Perl, D.P., Steiner, J., Pasternack, N., Li, W., Maric, D., Safavi, F., Horkayne-Szakaly, I., Jones, R., Stram, M.N., Moncur, J.T., Hefti, M., Folkerth, R.D., Nath, A., 2022. Neurovascular injury with complement activation and inflammation in COVID-19. Brain. https://doi.org/10.1093/brain/awac151

Lu, Y., Li, X., Geng, D., Mei, N., Wu, P.-Y., Huang, C.-C., Jia, T., Zhao, Y., Wang, D., Xiao, A., Yin, B., 2020a. Cerebral Micro-Structural Changes in COVID-19 Patients - An MRI-based 3-month Follow-up Study. EClinicalMedicine 25, 100484.

Lu, Y., Li, X., Geng, D., Mei, N., Wu, P.-Y., Huang, C.-C., Jia, T., Zhao, Y., Wang, D., Xiao, A., Yin, B., 2020b. Cerebral Micro-Structural Changes in COVID-19 Patients – An MRI-based 3-month Follow-up Study. EClinicalMedicine. https://doi.org/10.1016/j.eclinm.2020.100484

MacIntosh, B.J., Ji, X., Chen, J.J., Gilboa, A., Roudaia, E., Sekuler, A.B., Gao, F., Chad, J.A., Jegatheesan, A., Masellis, M., Goubran, M., Rabin, J., Lam, B., Cheng, I., Fowler, R., Heyn, C., Black, S.E., Graham, S.J., 2021. Brain structure and function in people recovering from COVID-19 after hospital discharge or self-isolation: a longitudinal observational study protocol. CMAJ Open 9, E1114–E1119.

Malerba, P., Agabiti Rosei, C., Nardin, M., Gaggero, A., Chiarini, G., Rossini, C., Fama’, F., Brami, V., Coschignano, M.A., Muiesan, M.L., Rizzoni, D., De Ciuceis, C., 2021. Early microvascular modifications in patients previously hospitalized for COVID-19: comparison with healthy individuals. Eur. Heart J. 42, ehab724.3387.

Matschke, J., Lütgehetmann, M., Hagel, C., Sperhake, J.P., Schröder, A.S., Edler, C., Mushumba, H., Fitzek, A., Allweiss, L., Dandri, M., Dottermusch, M., Heinemann, A., Pfefferle, S., Schwabenland, M., Sumner Magruder, D., Bonn, S., Prinz, M., Gerloff, C., Püschel, K., Krasemann, S., Aepfelbacher, M., Glatzel, M., 2020. Neuropathology of patients with COVID-19 in Germany: a post-mortem case series. Lancet Neurol. 19, 919–929.

McLarnon, J.G., 2021. A Leaky Blood–Brain Barrier to Fibrinogen Contributes to Oxidative Damage in Alzheimer’s Disease. Antioxidants. https://doi.org/10.3390/antiox11010102

Meinhardt, J., Radke, J., Dittmayer, C., Franz, J., Thomas, C., Mothes, R., Laue, M., Schneider, J., Brünink, S., Greuel, S., Lehmann, M., Hassan, O., Aschman, T., Schumann, E., Chua, R.L., Conrad, C., Eils, R., Stenzel, W., Windgassen, M., Rößler, L., Goebel, H.-H., Gelderblom, H.R., Martin, H., Nitsche, A., Schulz-Schaeffer, W.J., Hakroush, S., Winkler, M.S., Tampe, B., Scheibe, F., Körtvélyessy, P., Reinhold, D., Siegmund, B., Kühl, A.A., Elezkurtaj, S., Horst, D., Oesterhelweg, L., Tsokos, M., Ingold-Heppner, B., Stadelmann, C., Drosten, C., Corman, V.M., Radbruch, H., Heppner, F.L., 2021. Olfactory transmucosal SARS-CoV-2 invasion as a port of central nervous system entry in individuals with COVID-19. Nat. Neurosci. 24, 168– 175.

Moreno-Escobar, M.C., Feizi, P., Podury, S., Tandon, M., Munir, B., Alvi, M., Adcock, A., Sriwastava, S., 2021. Acute cerebellitis following SARS-CoV-2 infection: A case report and review of the literature. J. Med. Virol. 93, 6818–6821.

Pierpaoli, C., Basser, P.J., 1996. Toward a quantitative assessment of diffusion anisotropy. Magn. Reson. Med. 36, 893–906.

Reiken, S., Sittenfeld, L., Dridi, H., Liu, Y., Liu, X., Marks, A.R., 2022. Alzheimer’s-like signaling in brains of COVID-19 patients. Alzheimers. Dement. 18, 955–965.

Russo, M.V., McGavern, D.B., 2015. Immune Surveillance of the CNS following Infection and Injury. Trends in Immunology. https://doi.org/10.1016/j.it.2015.08.002

Ryu, J.K., McLarnon, J.G., 2009. A leaky blood-brain barrier, fibrinogen infiltration and microglial reactivity in inflamed Alzheimer’s disease brain. J. Cell. Mol. Med. 13, 2911–2925.

Serrano, G.E., Walker, J.E., Tremblay, C., Piras, I.S., Huentelman, M.J., Belden, C.M., Goldfarb, D., Shprecher, D., Atri, A., Adler, C.H., Shill, H.A., Driver-Dunckley, E., Mehta, S.H., Caselli, R., Woodruff, B.K., Haarer, C.F., Ruhlen, T., Torres, M., Nguyen, S., Schmitt, D., Rapscak, S.Z., Bime, C., Peters, J.L., Alevritis, E., Arce, R.A., Glass, M.J., Vargas, D., Sue, L.I., Intorcia, A.J., Nelson, C.M., Oliver, J., Russell, A., Suszczewicz, K.E., Borja, C.I., Cline, M.P., Hemmingsen, S.J., Qiji, S., Hobgood, H.M., Mizgerd, J.P., Sahoo, M.K., Zhang, H., Solis, D., Montine, T.J., Berry, G.J., Reiman, E.M., Röltgen, K., Boyd, S.D., Pinsky, B.A., Zehnder, J.L., Talbot, P., Desforges, M., DeTure, M., Dickson, D.W., Beach, T.G., 2022. SARS-CoV-2 Brain Regional Detection, Histopathology, Gene Expression, and Immunomodulatory Changes in Decedents with COVID-19. J. Neuropathol. Exp. Neurol. https://doi.org/10.1093/jnen/nlac056

Shaikh, A.G., Manto, M., Mitoma, H., 2022. 2 years into the pandemic: What did we learn about the COVID-19 and cerebellum? Cerebellum 21, 19–22.

Shankar, S.K., Mahadevan, A., Kovoor, J.M.E., 2008. Neuropathology of Viral Infections of the Central Nervous System. Neuroimaging Clinics of North America. https://doi.org/10.1016/j.nic.2007.12.009

Stefanou, M.-I., Palaiodimou, L., Bakola, E., Smyrnis, N., Papadopoulou, M., Paraskevas, G.P., Rizos, E., Boutati, E., Grigoriadis, N., Krogias, C., Giannopoulos, S., Tsiodras, S., Gaga, M., Tsivgoulis, G., 2022. Neurological manifestations of long-COVID syndrome: a narrative review. Ther. Adv. Chronic Dis. 13, 20406223221076890.

van den Enden, A.J.M., van Gils, L., Labout, J.A.M., van der Jagt, M., Moudrous, W., 2020. Fulminant cerebral edema as a lethal manifestation of COVID-19. Radiology Case Reports. https://doi.org/10.1016/j.radcr.2020.06.053

Wenzel, J., Schwaninger, M., 2022. How COVID-19 affects microvessels in the brain. Brain awac211.

Westman, J., Grinstein, S., Marques, P.E., 2019. Phagocytosis of Necrotic Debris at Sites of Injury and Inflammation. Front. Immunol. 10, 3030.

Wong, A., Glaister, J., Cameron, A., Haider, M., 2013. Correlated diffusion imaging. BMC Med. Imaging 13, 26.

Wong, A., Gunraj, H., Sivan, V., Haider, M.A., 2021. Synthetic Correlated Diffusion Imaging Hyperintensity Delineates Clinically Significant Prostate Cancer. arXiv [physics.med-ph].

Wong, A., Khalvati, F., Haider, M.A., 2015. Dual-stage correlated diffusion imaging, in: 2015 IEEE 12th International Symposium on Biomedical Imaging (ISBI). pp. 75–78.

Wright, P.W., Vaida, F.F., Fernández, R.J., Rutlin, J., Price, R.W., Lee, E., Peterson, J., Fuchs, D., Shimony, J.S., Robertson, K.R., Walter, R., Meyerhoff, D.J., Spudich, S., Ances, B.M., 2015. Cerebral white matter integrity during primary HIV infection. AIDS. https://doi.org/10.1097/qad.0000000000000560

Xydakis, M.S., Albers, M.W., Holbrook, E.H., Lyon, D.M., Shih, R.Y., Frasnelli, J.A., Pagenstecher, A., Kupke, A., Enquist, L.W., Perlman, S., 2021. Post-viral effects of COVID-19 in the olfactory system and their implications. Lancet Neurol. 20, 753–761.

