## Supplementary Materials for "Sensitivity of diffusion-tensor and correlated diffusion imaging to white-matter microstructural abnormalities: application in COVID-19"

### Theory

#### Diffusion tensor imaging

##### Conventional DTI

In the conventional DTI model, the k[th] signal (in terms of diffusion-gradient direction) in DTI is given by

$$S_k = S_0 e^{-b q_k^T D q_k} \quad \text{(A1)}$$

Where b is the b-value, **D** is the diffusion tensor, and $q_k$ is the unit vector for the $k^{\text{th}}$ diffusion direction. **D** is a second-order tensor (a matrix), is symmetric and describes 3D diffusion, so it is 3x3 matrix. DTI studies most commonly employ the metrics mean diffusivity (MD), a measure of the size of the tensor, and fractional anisotropy of diffusion (FA), a measure of the shape of the tensor, both derived from the diffusion tensor,

$$MD = \frac{1}{3} Tr[D] - mean(\lambda) \quad \text{(A2)}$$

$$FA = \sqrt{\frac{3}{2}} \frac{NA}{\sqrt{NA^2 + 3MD^2}} \quad \text{(A3)}$$

Where Tr is the trace of **D**, lambda is the set of eigenvalues for **D**, and NA is the norm of anisotropy, is a shape measure directly based on the variance of the eigenvalues and thus irrespective of alterations in MD (i.e. irrespective of alterations in isotropic diffusion).

$$NA = ||D - MD|| \quad \text{(A4)}$$

where $||\quad||$ denotes the Frobenius norm. While FA also describes the shape of the tensor, the measure is normalized by the size of the tensor, and hence FA and MD are not orthogonal. Furthermore, MD can be broken down into axial (AD) and radial diffusivity (RD), corresponding to the primary axis and the radial direction to the tensor, respectively,

$$AD = \lambda_1 \quad \text{(A5)}$$

$$RD = \frac{\lambda_2 + \lambda_3}{2} \quad \text{(A6)}$$

DT-DOME

To provide a more holistic representation of anisotropy rather than quantifying directional diffusivities individually, we can choose an alternative decomposition of the diffusion tensor. We recently proposed an alternative orthogonal decomposition based on the eigenvalue moments (DT-DOME): MD, NA and mode of anisotropy (MO) (Ennis and Kindlmann, 2006).

$$MO = 3\sqrt{6}det(\frac{D-MD}{||D-MD||}) \quad \text{(A7)}$$

where det represents the matrix determinant. Note that degeneration of primarily single-tract regions would manifest as decreased NA, whereas selective degeneration of secondary crossing tracts would manifest as increased NA. MO is another shape measure that corresponds with the skewness of the eigenvalues and hence whether the anisotropy is linear (MO=1), or planar (MO=-1). DT-DOME has demonstrated superior sensitivity to fibre shape compared to conventional DTI, especially in regions of complex fibre architectures which permeate the white matter (Chad et al., 2021).

#### Correlated diffusion imaging (CDI)

As shown in Eq. 1, the diffusion signal amplitude itself embodies information on tissue diffusivity. That is, restricted diffusion can manifest as increased diffusion-weighted signal intensity, and by combining this signal intensity over many acquisitions, the difference between different degrees of diffusion restriction could be enhanced. In the first demonstration of CDI, a diffusion-weighted data set was acquired by varying the diffusion-gradient strength (and not the gradient direction), and the CDI value for each voxel was defined as

$$C(x) = \int \ldots \int S_{q_\alpha}(x)\ldots S_{q_\alpha}(x) f(S_{q_\alpha}(x)\ldots S_{q_\beta}(x)|V(x) \times dS_{q_\alpha}(x)\ldots dS_{q_\beta}(x)$$

(A8)

where $q_\alpha$ and $q_\beta$ indicate the two ends of the range of diffusion weightings, $x$ indicates a given spatial location surrounded by a volume $V(x)$, $S$ represents the diffusion-weighted signal intensity, and $f$ represents the conditional joint probability density function. $V(x)$ should be chosen according to the scale of the expected abnormality. In cancer tissue, since tissue anisotropy is low, $f$ can be a symmetric 2D kernel. Intuitively, a higher CDI value translates to a lower MD, but CDI intensity demonstrated greater sensitivity and specificity than MD for detecting prostate cancer in over 200 patients (Wong et al., 2013). Thus, it was identified as a possible candidate for targeting subtle neuroinflammatory effects associated with COVID-19.

In the cerebral white matter, given the need to minimize partial-volume effects, $V(x)$ and $f$ should both be minimized in width (to 1 voxel). Also, diffusion MRI acquisitions typically span a range of gradient strengths and directions. Thus, the CDI intensity simplifies to

$$C(x) = \prod_{i=\alpha}^{\beta} \prod_{j=a}^{b} S_{q_{i,j}}(x) \quad \text{(A9)}$$

Where *i* and *j* are indices traversing the range of diffusion weightings (spanning $\alpha$ and $\beta$) and directions (spanning a and b), respectively.

#### Simulation results

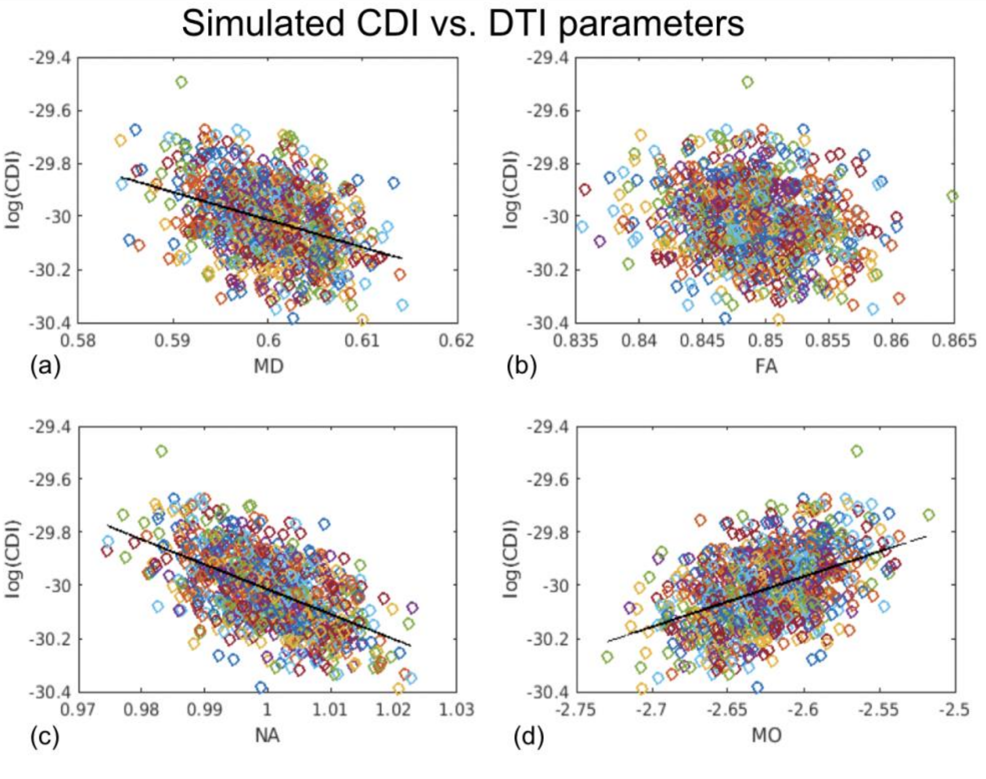


**Figure A1. Relationship between CDI and DTI parameters.** Values are plotted for the highest simulated SNR of 110. (a, c) log(CDI) is significantly inversely correlated with MD and NA; (b) there is no clear trend linking CDI and FA; (d) log(CDI) is significantly positive correlated with MO.

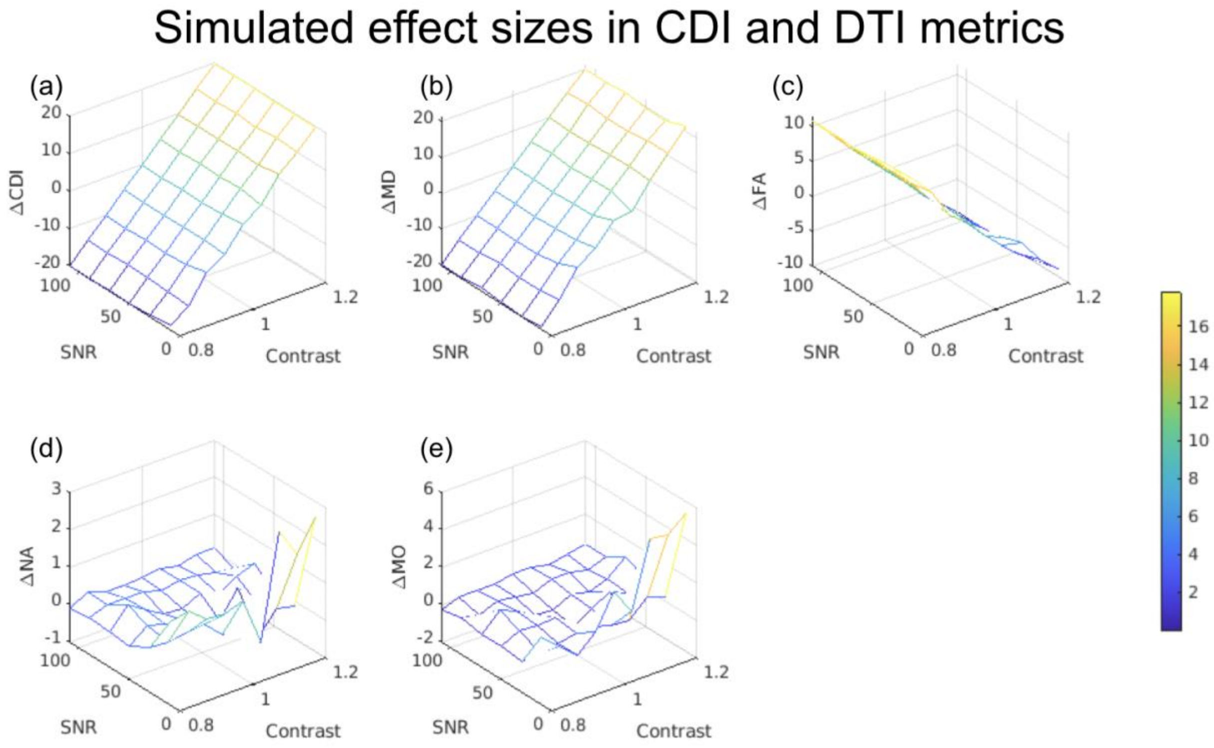


**Figure A2. Estimated effect sizes associated with CDI and DTI metrics for distinguishing healthy and diseased white-matter tissue.** Each vertex on the 3D mesh represents 1 data point. Effect size is given as a percentage indicated by the colourbar. The effect size is comparable between log(CDI) (a) and MD (b), much lower for FA (c), and lower still for NA (d) and MO (e).

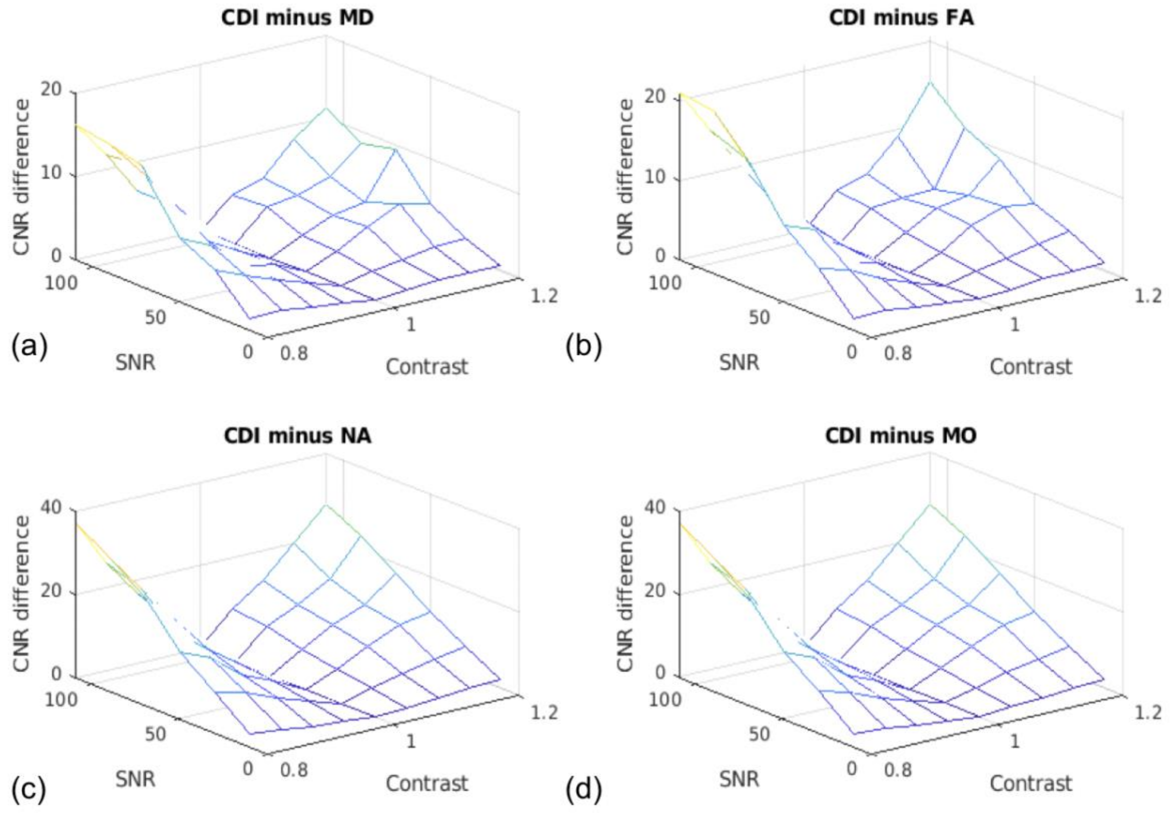


**Figure A3. CNR differences between log(CDI) and conventional-DTI metrics.** In all cases, log(CDI) conferred a CNR advantage when differentiating between healthy and diseased tissue (as indicated by large positive CNR differences). .

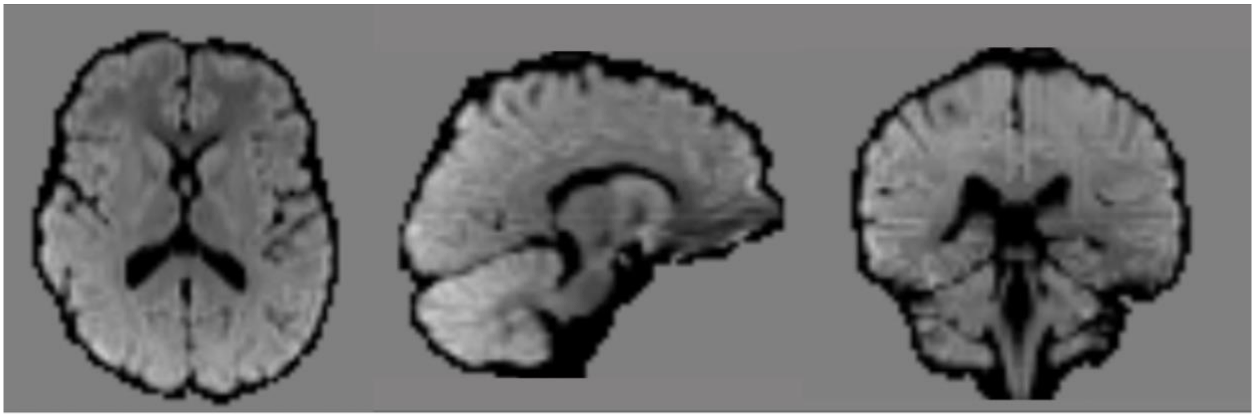

**Figure A2. Sample log(CDI) maps from a representative subject.**
